## Supplemental Figures for "Kinetic regulation of kinesin’s two motor domains coordinates its stepping along microtubules"

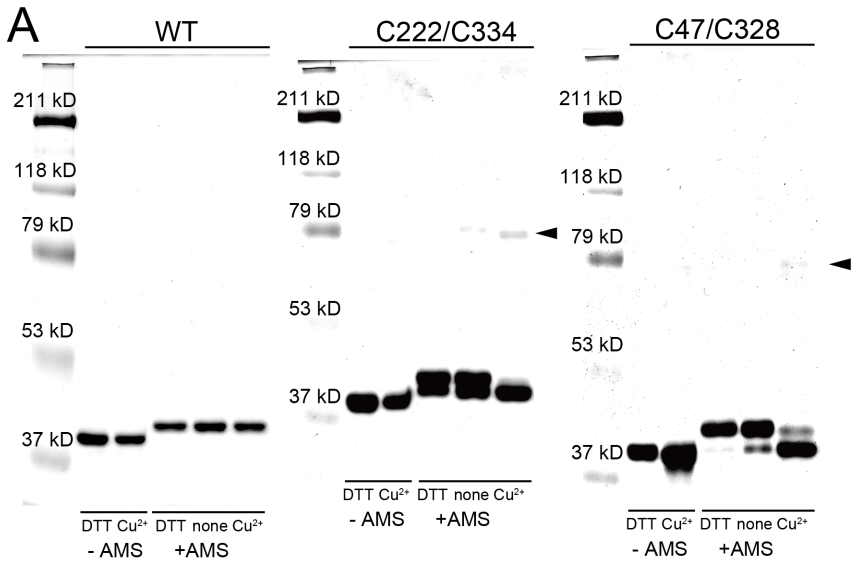


B

| | | Uncrosslinked | Intramolecular crosslinked | Intermolecular crosslinked |
|---|---|---|---|---|
| C222/C334 | DTT | 70.2 % | 29.8 % | 0 % |
| | None | 62 % | 37.7 % | 0.3 % |
| | $Cu^{2+}$ | 0 % | 97.3 % | 2.7 % |
| C47/C328 | DTT | 98.4 % | 1.6 % | 0 % |
| | None | 88.7 % | 11.3 % | 0 % |
| | $Cu^{2+}$ | 11.1 % | 86.8 % | 2.1 % |

**Figure 1–figure supplement 1.** Estimation of disulfide-crosslinking efficiency. (**A**) The whole gel image of the SDS-PAGE under non-reducing conditions with enhanced contrast to highlight the minor bands (close up of these gels are shown in **Figure 1D**). Cys-light K339 construct with no cysteine introduction (WT), with Cys222/Cys334 and with Cys47/Cys328 were treated with either reducing (10 mM DTT), oxidative (10 µM $Cu^{2+}$ and 20 µM o-phenanthroline) conditions, or none (neither DTT nor $Cu^{2+}$). 20 mM AMS, which covalently modifies free thiol residues, was added to enhance the band shift after crosslinking. Arrowheads indicate higher molecular weight bands (double the molecular weight of a monomer), corresponding to intermolecular disulfide cross-linked molecules. Cys-light K339 kinesin without cysteine introduction (WT) showed no band shift after oxidative treatment, confirming that crosslinking occurred only between substituted residues. (**B**) Ratio of the un-crosslinked, disulfide intramolecular-crosslinked, and disulfide intermolecular-crosslinked bands, as estimated from the intensities of the bands. Under oxidative conditions, un-crosslinked and intermolecular-crosslinked species were minor and would not significantly affect the rate constants of the bulk measurements.

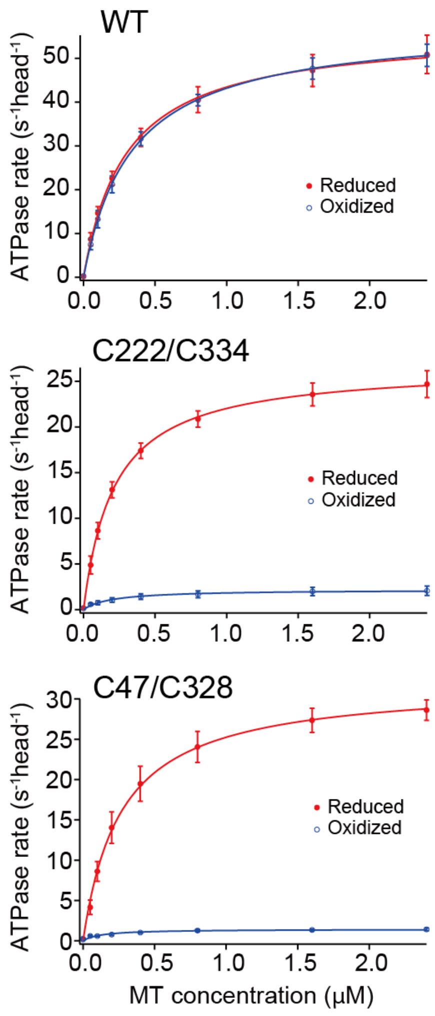


**Figure 1–figure supplement 2.** Steady-state microtubule-activated ATPase rates of monomeric kinesins before and after oxidative crosslinking. These rates, measured under reducing and oxidizing conditions, were plotted against microtubule (MT) concentrations for monomeric kinesins with and without cysteine substitutions (N = 3; average ± s.e.m.). The solid lines show fit with the Michaelis-Menten equation, and the fit parameters ($k_{cat}$ and $K_m$(MT)) are summarized in **Table 1**.

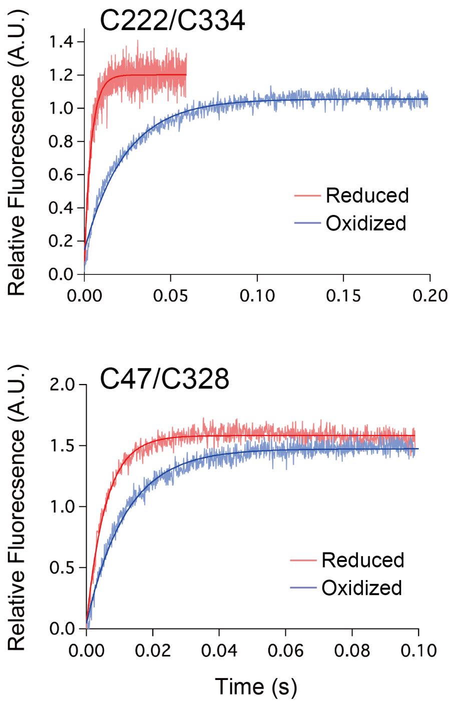


**Figure 2–figure supplement 1.** Pre-steady-state kinetics of ATP binding to monomeric kinesin-microtubule complex before and after crosslinking. Examples of fluorescence signal time trace for C222/C334 and C47/C328 monomers bound to the microtubule under reduced (red) and oxidized (blue) conditions after rapid mixing with 20 μM mant-ATP. The solid lines represent the fit with a single exponential to determine the observed rate constants ($k_{obs}$).

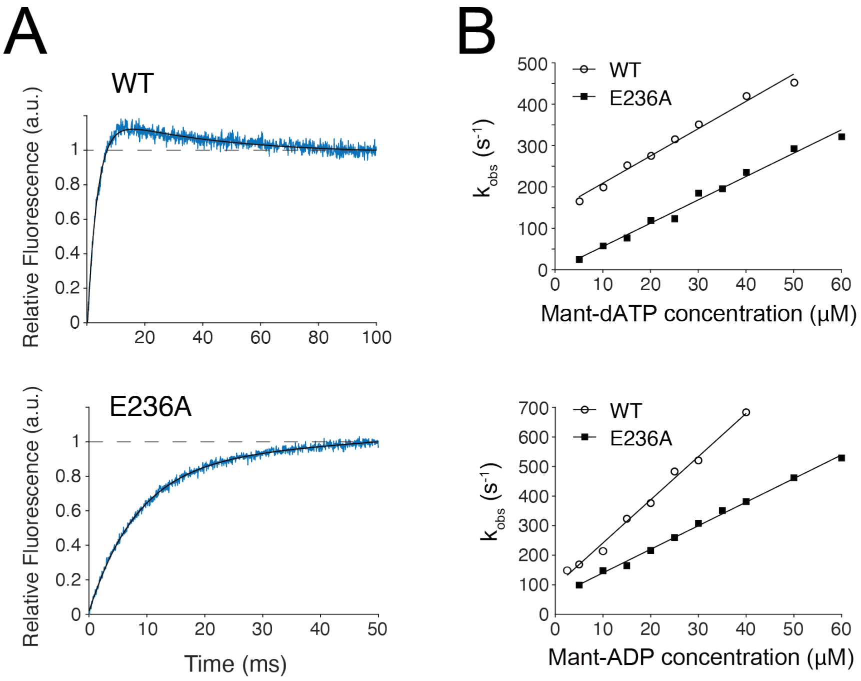


**Figure 2–figure supplement 2.** ATP binding kinetics for E236A mutant monomer on microtubule. (**A**) Examples of fluorescence signal time trace for microtubule-bound wild-type (WT) and E236A monomers (K349 without Cys-substitution) after rapid mixing with 20 μM mant-dATP. The WT monomer traces were fitted to a double exponential due to the presence of a decrease phase, previously reported by Ma and Taylor (1997a). The E236A monomer traces were fitted to a burst equation (Eq. 2 in Materials and Methods). (**B**) The $k_{obs}$ plot as a function of mant-dATP or mant-ADP concentrations. Solid lines represent a linear fit. The fit parameters for mant-dATP binding were: $k_{+1}$ = 6.6 ± 0.3 $\mu M^{-1}s^{-1}$ and $k_{-1}$ = 143 ± 9 $s^{-1}$ for wild-type monomer, and $k_{+1}$ = 5.6 ± 0.2 $\mu M^{-1}s^{-1}$ and $k_{-1}$ = −0.6 ± 7.7 $s^{-1}$ for E236A monomer. For mant-ADP binding: $k_{+1}$ = 14.6 ± 0.5 $\mu M^{-1}s^{-1}$ and $k_{-1}$ = 95 ± 12 $s^{-1}$ for wild-type monomer, and $k_{+1}$ = 8.0 ± 0.2 $\mu M^{-1}s^{-1}$ and $k_{-1}$ = 61 ± 6 $s^{-1}$ for E236A monomer. E236A showed a reduced ATP off-rate ($k_{-1}$) while maintaining an ADP off-rate comparable to wild-type.

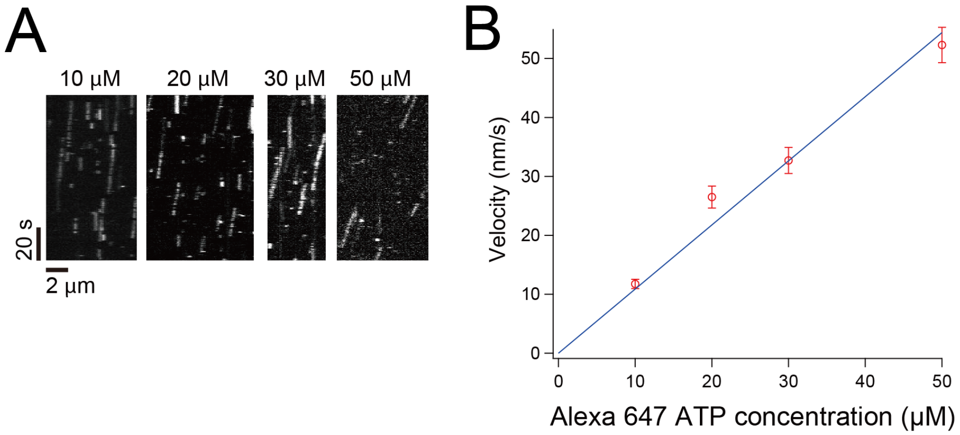


**Figure 2–figure supplement 3.** Processive motility of wild-type kinesin dimer driven by Alexa 647 ATP. (**A**) Kymographs showing the motilities of the ATTO 488-labeled K490 dimeric kinesin in the presence of varying amounts of Alexa 647-conjugated ATP, recorded at 5 fps. (**B**) Mean velocities of the processive movement were plotted against Alexa 647-ATP concentrations. The solid line represents a fit with a linear line under the fixed zero intercept; the slope was 1.09 ± 0.05 nm/s/μM. The velocities were about 10-fold smaller than those observed in the presence of unmodified ATP (495 nm/s at 50 μM ATP; Isojima et al., 2016), suggesting that the Alexa-conjugated ATP binds to the head with lower affinity than unmodified ATP.

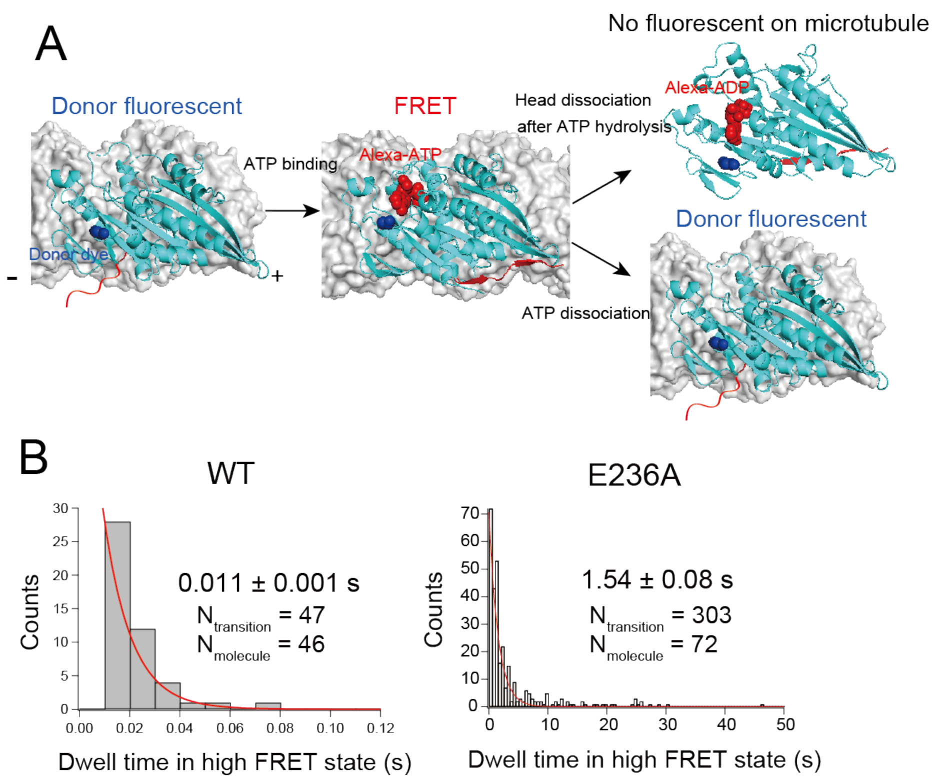


**Figure 2–figure supplement 4.** Dwell time of Alexa-ATP-bound state for wild-type and E236A monomers determined using smFRET. (**A**) Schematic showing the single-molecule FRET observation of ATP binding to the monomer head, observed using total-internal reflection fluorescent microscopy. The Cys55 (blue sphere) on Cys-light K349 monomeric kinesin, which is located close to the nucleotide pocket, was labeled with a donor fluorescent dye, and observed in the presence of the acceptor-labeled ATP (Alexa-ATP; red). The FRET efficiency increases up to ~90% when Alexa-ATP binds to the donor-labeled head on the microtubule. The donor and acceptor fluorescent both disappear when the head hydrolyses ATP and detaches from the microtubule. Conversely, the donor fluorescent recovers when the bound Alexa-ATP dissociates from the head reversibly. (**B**) Single-molecule FRET between ATTO488-labeled wild-type or Cy3-labeled E236A monomeric kinesin and Alexa 647 ATP was observed. Typical time traces are shown in **Figure 2D, E**. Histograms of the dwell time of the high FRET state were fit with a single exponential (red lines). The numbers represent the average dwell times (± s.e.m.) determined from the fit. The high FRET state of the wild-type monomer includes head dissociation after ATP hydrolysis and reversible ATP dissociation. The inverse of the mean dwell time for wild-type and E236A monomers are 91.5 ± 4.7 $s^{-1}$ and 0.65 ± 0.04 $s^{-1}$, respectively (**Figure 2F**). The inverse of the mean dwell time for head dissociation and ATP dissociation for the wild-type were 89.7 ± 6.7 $s^{-1}$ (N=26) and 93.5 ± 9.4 $s^{-1}$ (N=20), respectively; they are nearly indistinguishable, in part because of limited temporal resolution (10 ms).

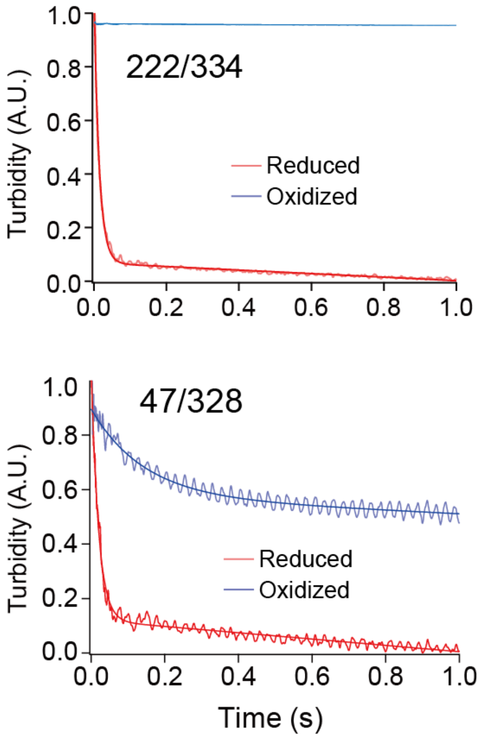


**Figure 3–figure supplement 1.** ATP-induced dissociation kinetics of crosslinked kinesin from microtubule measured using light scattering. Turbidity (absorption at 340 nm) was monitored after rapidly mixing the kinesin-microtubule complex with 1 mM ATP and 150 mM KCl. The solid lines represent a fit to a burst equation (Eq. 2 in Materials and methods). To prevent the motors from rebinding to the microtubule, we increased the KCl concentration of the kinesin-microtubule solution to 150 mM immediately before loading into the syringe. The C222/C334 did not show any initial rapid decrease in turbidity, presumably because the crosslinked C222/C334 has low microtubule-affinity and would immediately dissociate from the microtubule after increasing the KCl concentration.

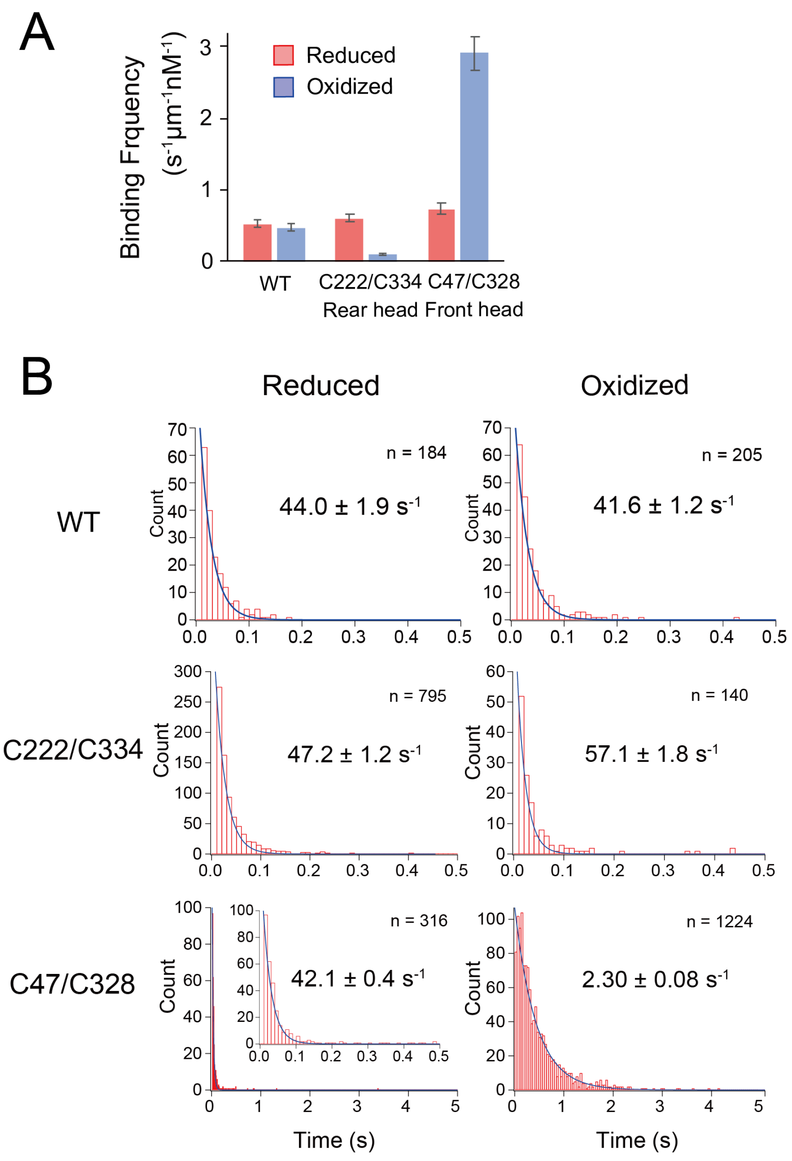


**Figure 3–figure supplement 2.** ATP-dependent microtubule-binding and -dissociation of crosslinked kinesins observed by single-molecule fluorescence microscopy. (**A**) Frequency of binding of GFP-labeled kinesin to microtubule. The binding/unbinding of GFP-fused kinesin to/from microtubule was observed in the presence of 1 mM ATP and 100 mM KCl using total-internal reflection fluorescent microscopy (refer to **Figure 3B** for kymographs). Number of fluorescent spots were divided by time period (s), microtubule length (μm) and kinesin concentration (nM). The frequency decreased six-fold for C222/C328 and increased four-fold for C47/C328, demonstrating that forward and backward constraints alter the microtubule affinity in different ways. (**B**) Histograms of the dwell time of GFP-labeled kinesin on microtubule. Solid lines represent a fit to an exponential decay curve. The numbers represent the average dwell times (± s.e.m.) determined from the fit.

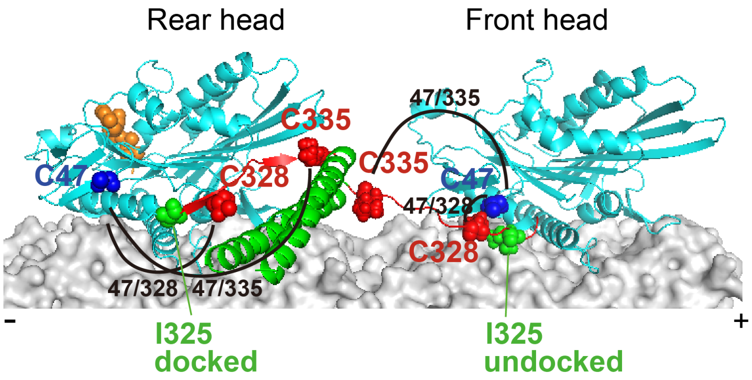


**Figure 3–figure supplement 3.** Locations of the C47/C328 and C47/C335 residues when the labeled head is in the front or rear of dimeric kinesin. The crosslinking between C47 (blue sphere) and C328 (red sphere) prevents the I325 residue (green sphere) from reaching the hydrophobic pocket on the head, as depicted in the front head. On the other hand, the crosslinking of C47 and C335 (red sphere) creates a flexible loop, which allows the initial segment of the neck linker, which includes the I325 residue, to dock onto the head as shown in the rear head.

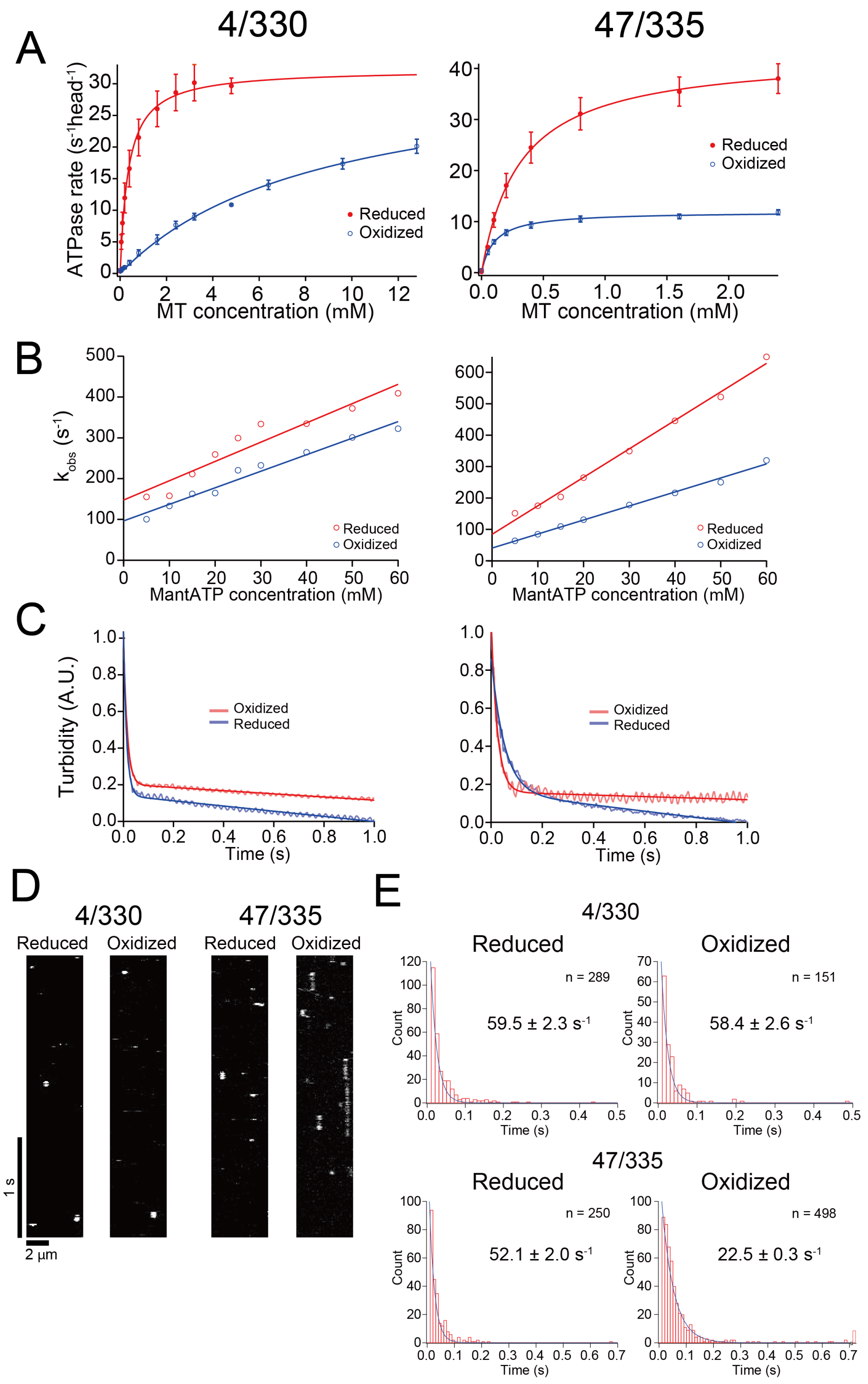

A
4/330
47/335
ATPase rate (s-1head-1)
MT concentration (mM)
Reduced
Oxidized
B
kobs (s-1)
MantATP concentration (mM)
Reduced
Oxidized
C
Turbidity (A.U.)
Time (s)
Oxidized
Reduced
D
4/330
47/335
Reduced
Oxidized
Reduced
Oxidized
1 s
2 µm
E
4/330
Reduced
Oxidized
n = 289
59.5 ± 2.3 s-1
n = 151
58.4 ± 2.6 s-1
47/335
Reduced
Oxidized
n = 250
52.1 ± 2.0 s-1
n = 498
22.5 ± 0.3 s-1
Count
Time (s)

**Figure 3–figure supplement 4.** Kinetics measurements of ATP-binding and microtubule-dissociation of C4/C330 and C47/C335 crosslinked monomeric kinesins. Positions of cysteine residues for disulfide crosslinking are shown in **Figure 3D**. The crosslinking efficiencies estimated by SDS-PAGE under non-reducing conditions were 92 and 85% for 4/330 and 47/335, respectively. Steady-state microtubule-activated ATPase rates (**A**), pre-steady-state mant-ATP binding kinetics (**B**), pre-steady-state microtubule-dissociation kinetics (**C**), kymographs of single-molecule fluorescence microscopy of GFP-fused monomers with 1 mM ATP and 100 mM KCl (**D**), and histograms of dwell time on microtubule (**E**), before and after oxidative crosslinking. The mean values and fit parameters are summarized in **Table 1**

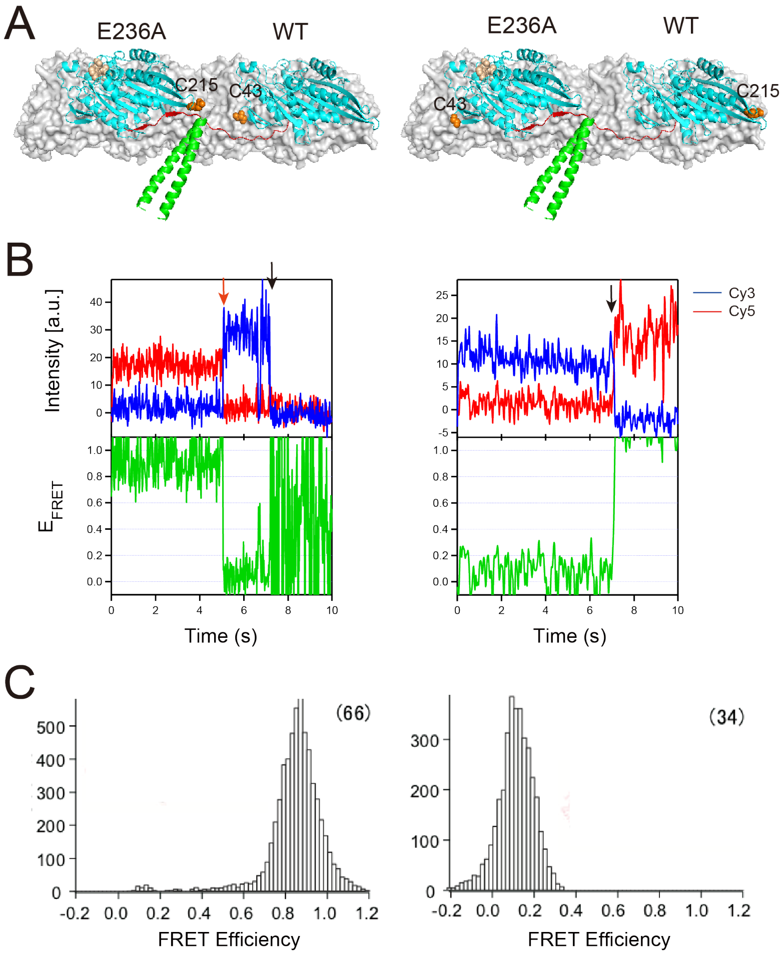


**Figure 4–figure supplement 1.** Single-molecule FRET between two heads of E236A-WT heterodimer. (**A**) Schematic diagrams showing the positions of introduced cysteines (orange spheres) for labeling with donor/acceptor fluorophores; left; 215Cys on E236A-chain and 43Cys on WT-chain, right; 43Cys on E236A-chain and 215Cys on WT-chain. When the E236A head and WT head occupy the trailing and leading positions, respectively, the distances between these two cysteine residues are ~3 nm and ~13 nm. (**B**) Typical traces of fluorescence intensities of donor (Cy3, blue) and acceptor (Cy5, red) fluorophores, as well as the calculated FRET efficiency ($E_{FRET}$), for 215-43 labeled E236A-WT heterodimers, in the presence of 200 nM ATP at 50 fps. The red arrow indicates the photobleaching of the acceptor dye, while the black arrows indicate the photobleaching of the donor dye. We were unable to detect a transient FRET efficiency change associated with head detachment (toward ~30% FRET, or the one-head-bound state) at this temporal resolution, which requires measurements with a higher temporal resolution (**Figure 4**). (**C**) Histograms of FRET efficiencies (for each frame of images) of donor and acceptor labeled E236A-WT heterodimer, observed in the presence of 200 nM ATP. The number of molecules analyzed is shown in parentheses.

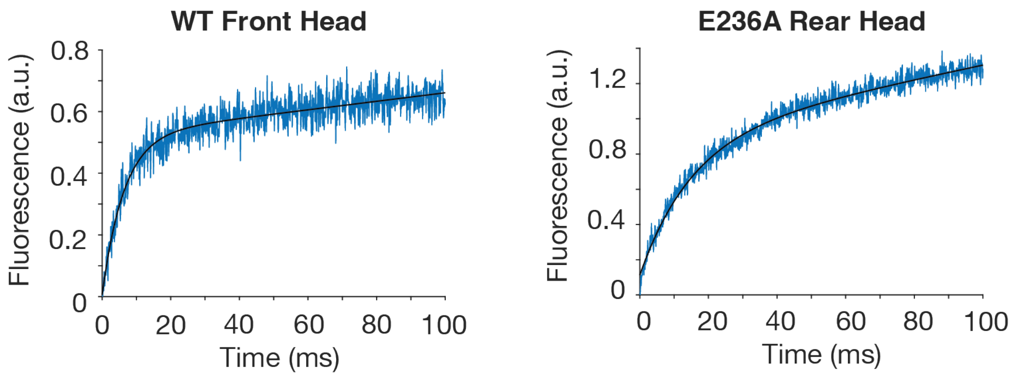


**Figure 4–figure supplement 2.** ATP binding kinetics for front and rear heads of E236A-WT heterodimer. Examples of fluorescence signal time trace for WT front head and E236A rear heads of E236A-WT heterodimer after rapid mixing with 15 μM mant-dATP. The traces were fitted to a burst equation. The $k_{obs}$ plots are shown in **Figure 4B**.

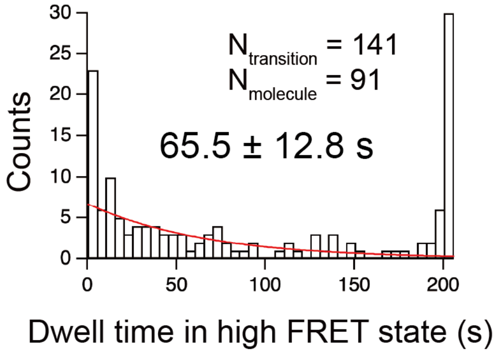


**Figure 4–figure supplement 3.** Single-molecule FRET observation between donor dye on the E236A head of E236A-WT heterodimer and acceptor-labeled ATP **(Figure 4D)**. Histogram of the dwell time of the high FRET state is shown. Typical time trace of the FRET efficiency is shown in **Figure 4E**. The solid lines show fit with an exponential decay (red lines). The number represents the average dwell times (± s.e.m.) determined from the fit. The inverse of the mean dwell time $k_{-1}$ is 0.015 ± 0.003 $s^{-1}$ **(Figure 4F)**.

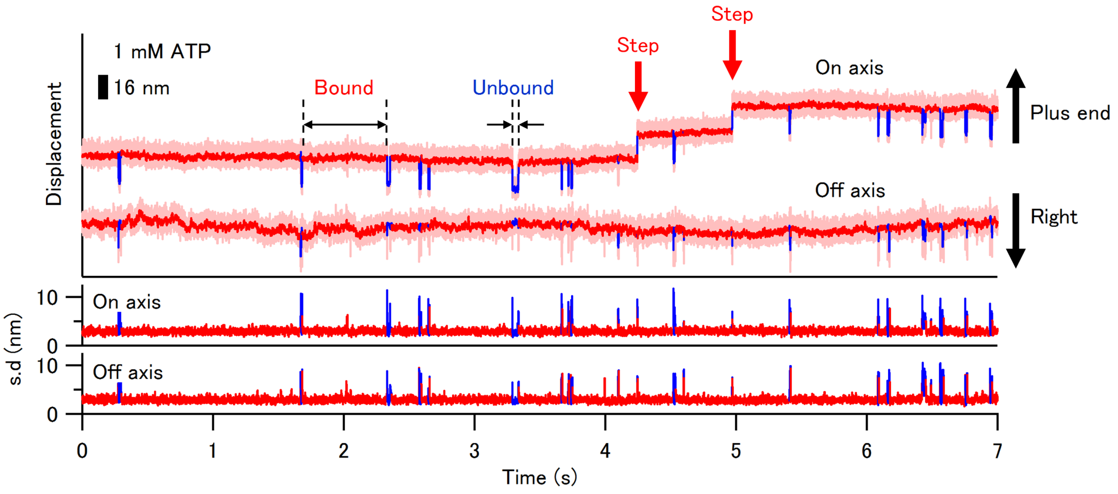


**Figure 5–figure supplement 1.** Long-term trace of the gold-labeled E236A-WT heterodimer exhibiting very slow stepping motion. A typical trace of the gold-labeled WT head of the E236A-WT heterodimer in the presence of 1 mM ATP recorded at 20,000 fps for 7 sec. The detached WT head mostly rebound to the same tubulin-binding site without exhibiting a step. However, the head occasionally bound to the 16-nm forward binding site (indicated by red arrows; forward-step), which happened about once every 10 sec (similar to the ATP hydrolysis rate of the E236A head). Upon unbinding, the detached WT head displaced toward the plus-end of the microtubule before stepping forward, which suggests that the WT head was in the trailing position before detachment. On the other hand, in the case of the rebinding, the detached head displaces toward the minus end of the microtubule, suggesting that the WT head was in the leading position before it detaches (refer to **Figure 5A**). If the trailing head detaches and rebinds to the same binding site, the displacement upon unbinding would be toward the plus-end (as seen in Fig. 5a in Isojima et al. 2016).

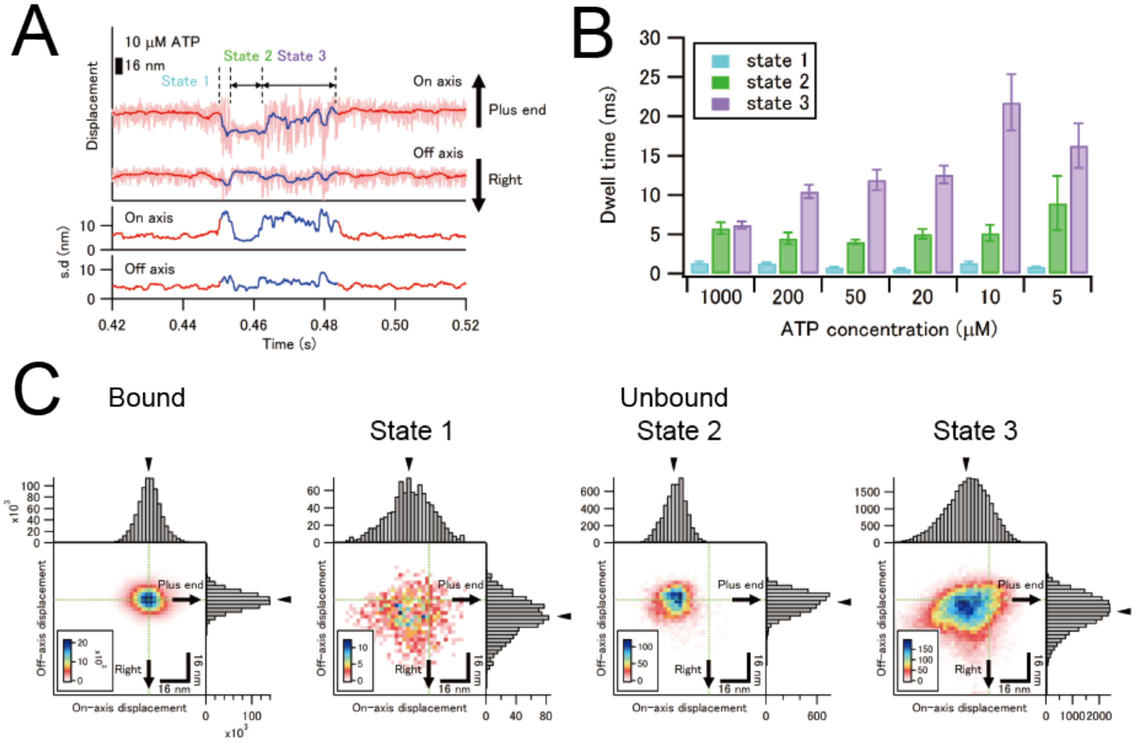


**Figure 5–figure supplement 2.** The unbound state of the gold-labeled WT head of the E236A-WT heterodimer. (**A**) A close-up view of the unbound state in the typical trace of gold-labeled WT head observed in the presence of 10 µM ATP (**Figure 5D**). This state is interrupted by a state of reduced fluctuation, which was not observed in the "trailing" head of the wild-type dimer as reported in Isojima et al. (2016). We refer to the initial high standard deviation (s.d.), second low s.d., and the last high s.d. states as state 1, 2, and 3. (**B**) The duration of each state in the unbound state. The dwell times for state 1 and 2 remain independent of ATP concentration (1.1 and 5.6 ms on average, respectively), whereas the dwell time for state 3 weakly depends on ATP concentration and is longer than those of state 1 and 2. (**C**) The ensemble-averaged 2-D histograms of the bound and unbound states of the gold-labeled WT head. Green lines for unbound states display the average on- and off-axis positions during the preceding bound state (representing the mean positions for the bound state). Arrowheads indicate the average position for the projected 1-D distributions. The fluctuation in the position of unbound state 2 is suppressed, much like the bound state, suggesting that the motility of the detached head is temporarily restrained. In contrast, the unbound state of G7 and G12 mutants with flexible insertions did not display the initial confined state (**Figure 6C**), suggesting that the confinement does not result from direct interaction between heads or between the head and the microtubule, but presumably rather from confinement on the neck linker of the unbound head. We propose that the detached front head initially exists in the ADP-bound/closed state (Sindelar et al., 2002; **Figure 7**), and its neck linker docks onto the head after microtubule-detachment (transitioning from state 1 to

2). In this state, both neck linkers are docked onto the head, and as a result, the mobility of the unbound head is limited due to the restricted degree of freedom in the neck linkers. Finally, the neck linker of the unbound head undocks, transitioning into the highly mobile state 3 (ADP-bound/semi-open state), which allows the unbound head to bind to the tubulin-binding site and release ADP.

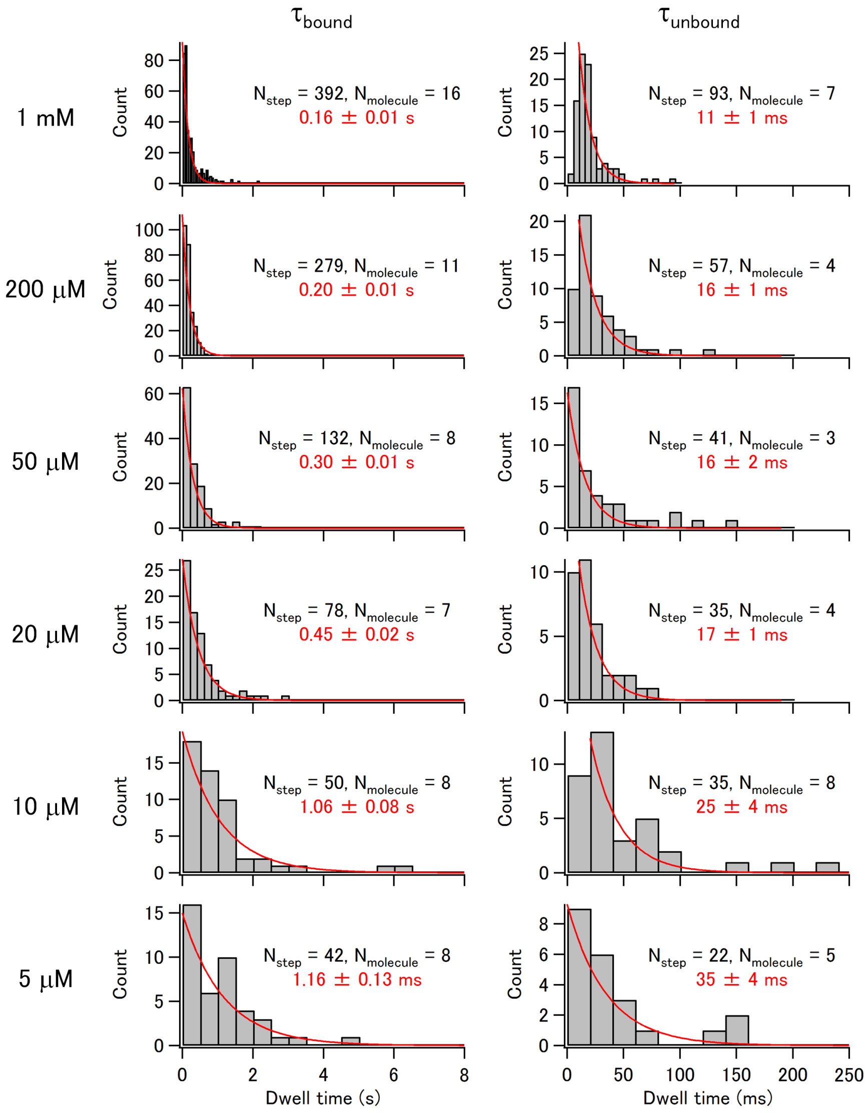


**Figure 5–figure supplement 3.** Distributions of the dwell time in the bound and unbound states of the leading WT head of E236A-WT heterodimer. Histograms of the dwell time in the bound ($\tau_{\text{bound}}$) and unbound ($\tau_{\text{unbound}}$) states of the gold-labeled WT head of the E236A-WT heterodimer at various ATP concentrations. Red lines indicate the fit with exponential decay. The average dwell times (± s.e.m.), as determined from the fits, are shown in red numbers and those of the bound state are summarized in **Figure 5E**.

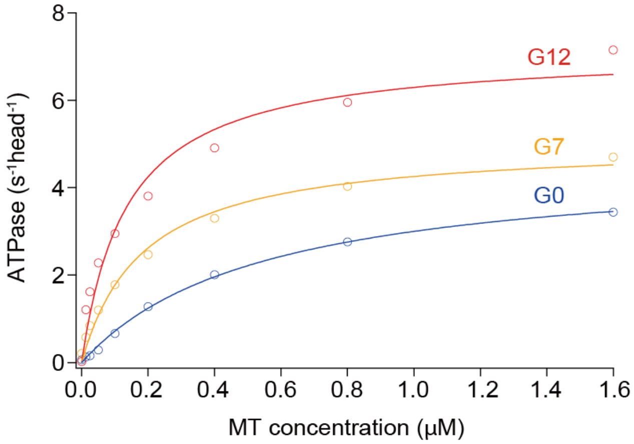


**Figure 6–figure supplement 1.** Microtubule-activated ATPase rates of the E236A-WT heterodimer with and without neck-linker extension. ATPase rates measured for E236A-WT heterodimer without neck-linker extension (termed G0), and with neck-linker extension of 7 (G7) or 12 poly-Gly insertion (G12) were plotted against microtubule (MT) concentrations. The solid lines represent the fit with the Michaelis-Menten equation. The fit parameters are $k_{cat}$ = 4.6 ± 0.1 ATP/s per head and $K_m$ (MT) = 0.54 ± 0.04 μM for G0, $k_{cat}$ = 5.0 ± 0.2 ATP/s per head and $K_m$ (MT) = 0.18 ± 0.03 μM for G7, and $k_{cat}$ = 7.2 ± 0.5 ATP/s per head and $K_m$ (MT) = 0.14 ± 0.03 μM for G12.

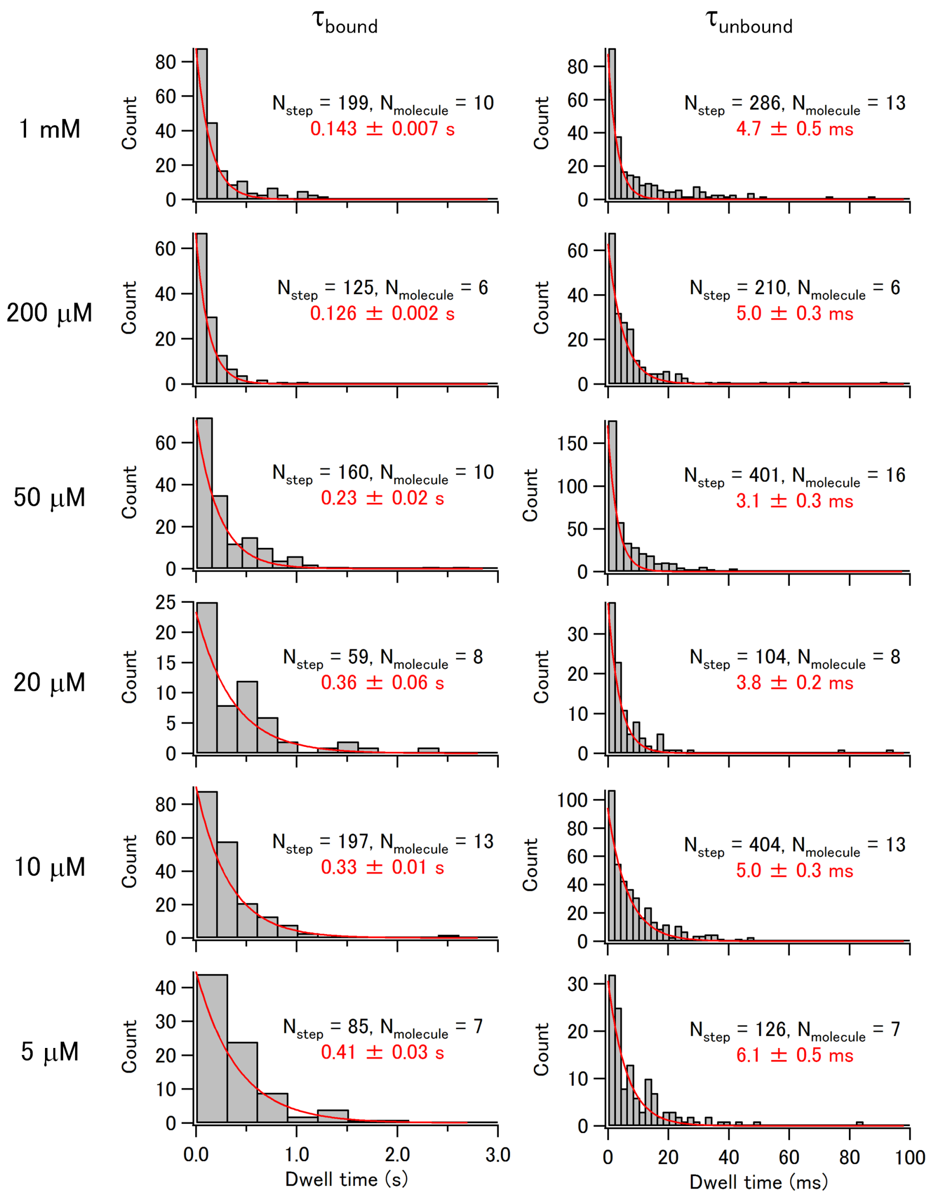


**Figure 6–figure supplement 2.** Distributions of the dwell time in the bound and unbound states of the leading WT head of E236A-WT heterodimer with 7 poly-Gly insertion (G7). Histograms of the dwell time in the bound ($\tau_{bound}$) and unbound ($\tau_{unbound}$) states of the gold-labeled WT head of the E236A-WT heterodimer at various ATP concentrations. Red lines indicate the fit with exponential decay. The average dwell times (± s.e.m.), as determined from the fits, are shown in red numbers. The inverse of the mean dwell time in the bound state $k_2$ is summarized in **Figure 6D**.

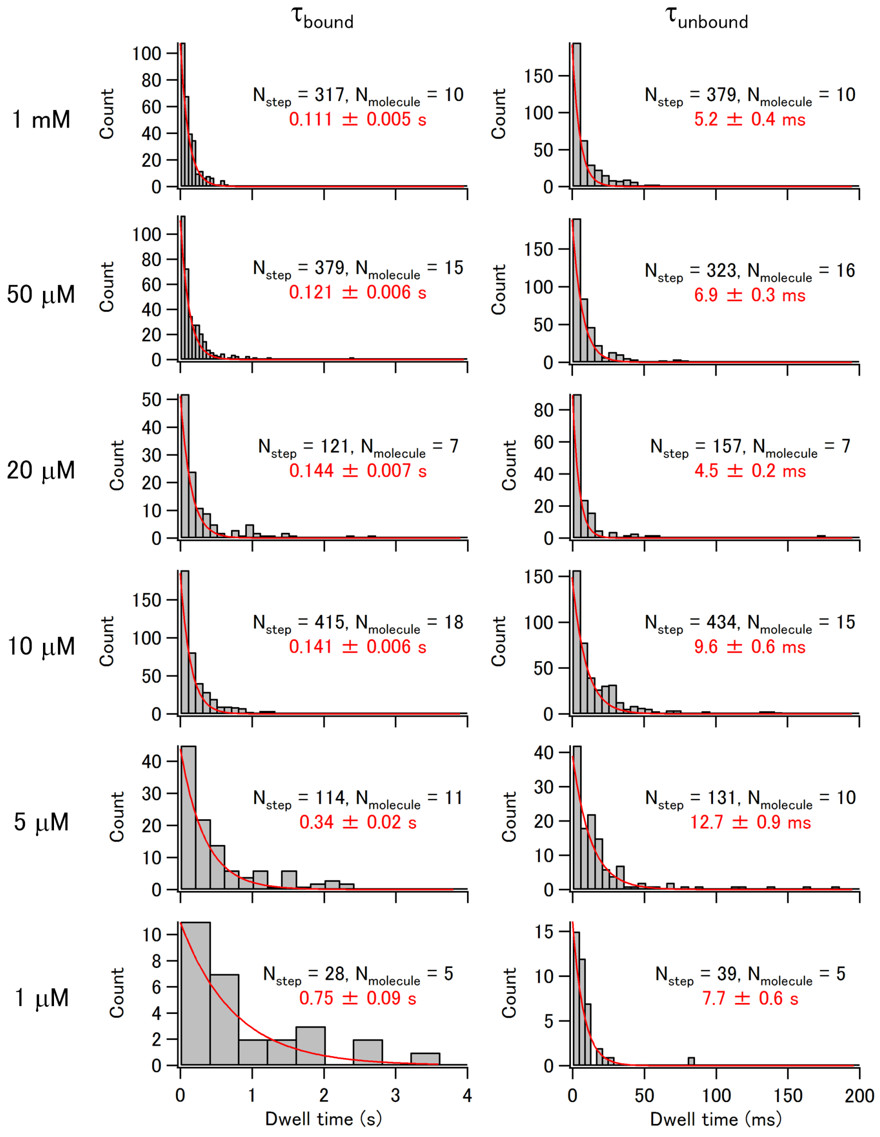


**Figure 6–figure supplement 3.** Distributions of the dwell time in the bound and unbound states of the leading WT head of E236A-WT heterodimer with 12 poly-Gly insertion (G12). Histograms of the dwell time in the bound ($\tau_{\text{bound}}$) and unbound ($\tau_{\text{unbound}}$) states of the gold-labeled WT head of the E236A-WT heterodimer at various ATP concentrations. Red lines indicate the fit with exponential decay. The average dwell times (± s.e.m.), as determined from the fits, are shown in red numbers. The inverse of the mean dwell time in the bound state $k_2$ is summarized in **Figure 6D**.

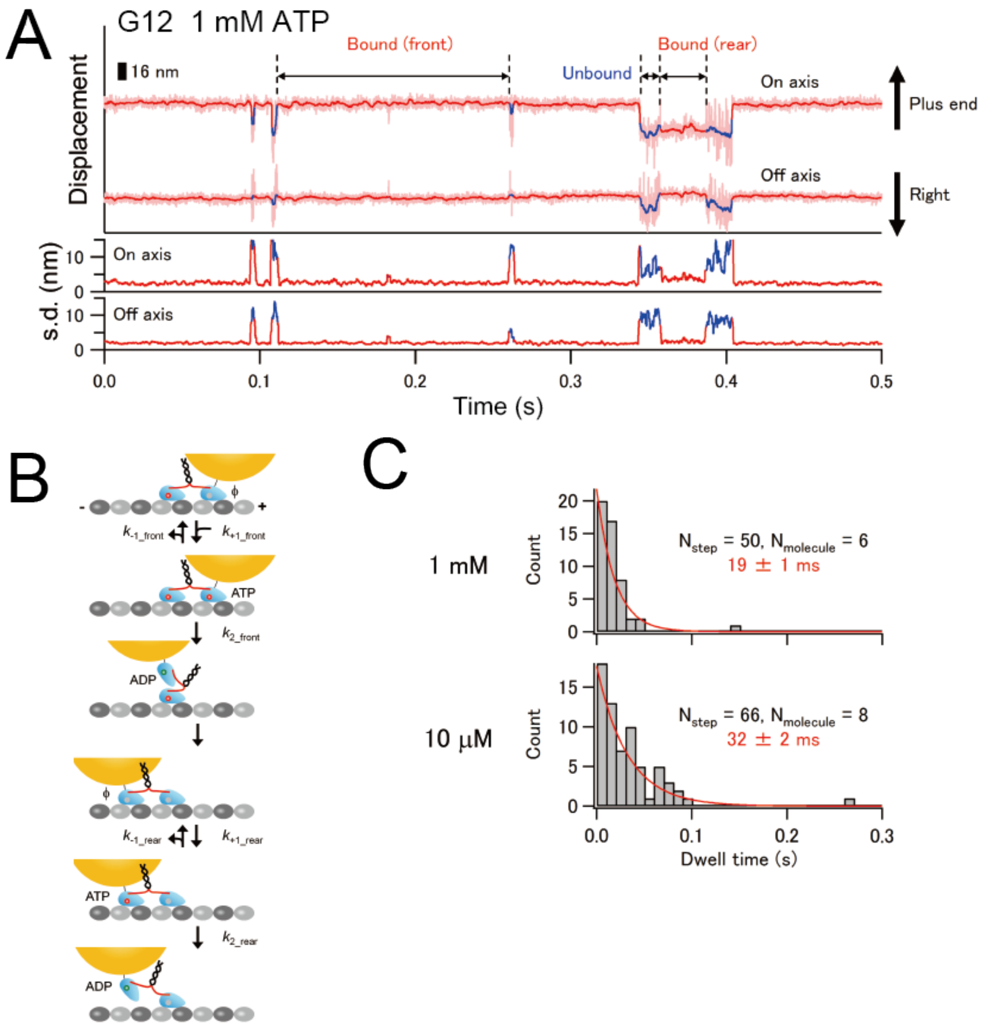


**Figure 6–figure supplement 4.** Binding to the rear-tubulin binding site observed for the G12 E236-WT heterodimer. (**A**) Typical trace for the centroid positions of the gold probe attached to the WT head of the E236A-WT heterodimer with a 12 poly-Gly insertion in the presence of 1 mM ATP. The WT head of the G12 heterodimer occasionally takes a backward step (i.e., binding to the rear-binding site 16-nm toward the minus-end of the microtubule; indicated as "bound (rear)"). The frequency of this backward step was 14% (N = 50 among 317 unbinding events) under the 1 mM ATP condition. (**B**) The schematic showing the observed backward step for the G12 heterodimer; the gold-labeled leading WT head detaches and binds to the rear-tubulin binding site. The dwell time of the bound state in the rear-binding site includes the ATP-binding rate $k_{+1\_\mathrm{rear}}$ and the ATP-promoted detachment rate $k_{2\_\mathrm{rear}}$. (**C**) Histograms of the dwell time of the bound state in the rear-binding site in the presence of 1 mM and 10 μM ATP. The dwell time under the 10 μM ATP condition was greater than that under the 1 mM ATP condition, suggesting that the bound state involves ATP-binding and -hydrolysis of the gold-labeled head.

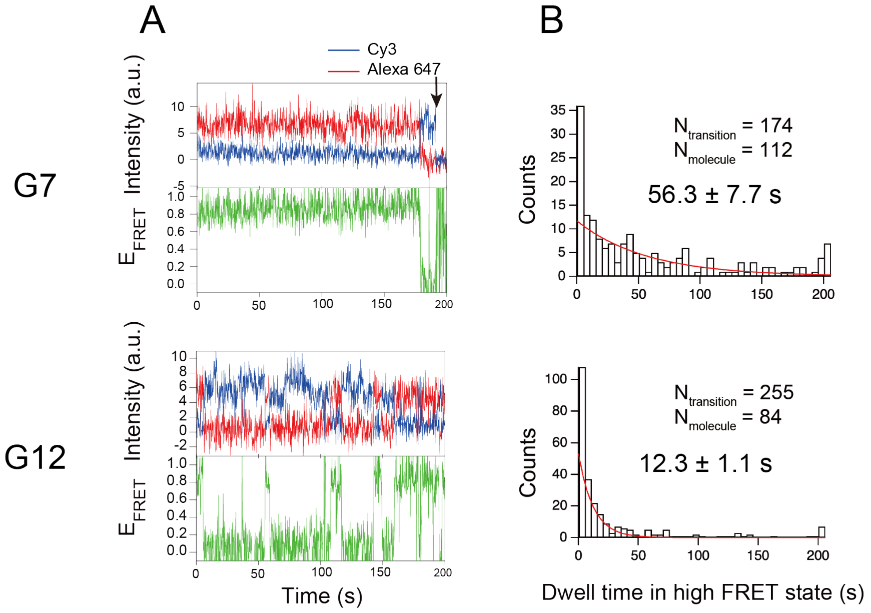


**Figure 6–figure supplement 5.** Single molecule FRET between donor-labeled E236A head of E236A-WT heterodimer with extended neck linker and acceptor-conjugated ATP. (**A**) Typical traces of fluorescence intensities of donor (Cy3, blue) and acceptor (Alexa 647, red) fluorophores, calculated FRET efficiency ($E_{\mathrm{FRET}}$), for E236A-WT heterodimer with Cy3 dye on the E236A head in the presence of 200 nM Alexa 647-conjugated ATP recorded at 5 fps. G7 and G12 represents E236A-WT heterodimer with 7 or 12 poly-Gly insertion between the neck linker and the neck coiled-coil. Black arrow indicates photobleaching of donor dye. (**B**) Histogram of the dwell time of high FRET state. The solid lines show fit with exponential decay and the mean dwell time (± s.e.m.) is shown within the histogram. The inverse of the mean dwell time for G7 and G12 E236A-WT heterodimer are 0.018 ± 0.002 $s^{-1}$ and 0.081 ± 0.006 $s^{-1}$, respectively **(Figure 5D)**.

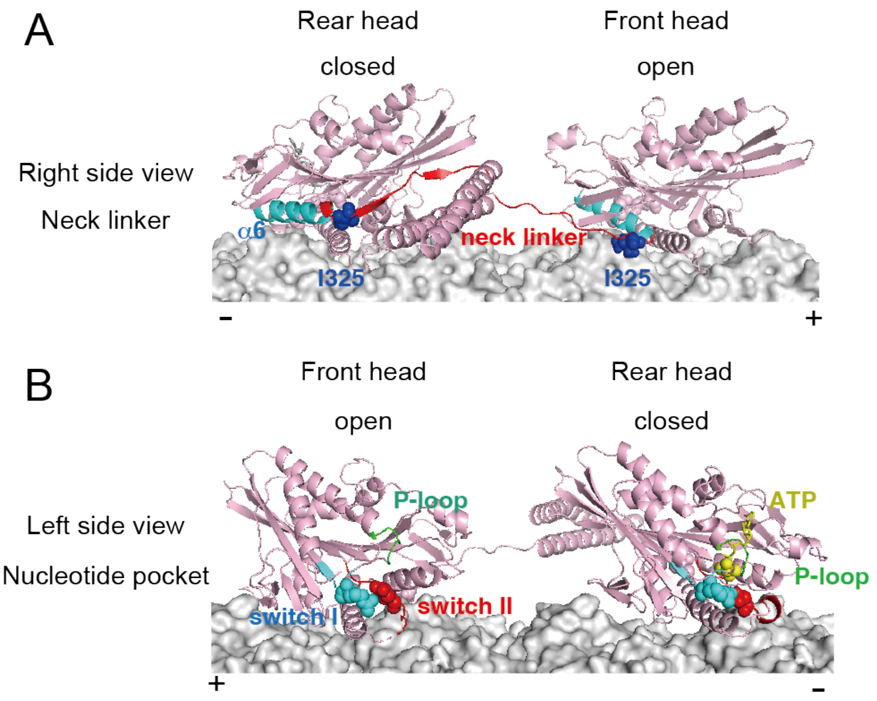


**Figure 7–figure supplement 1.** Structural difference between the open and closed conformational states of the kinesin head. The structural model of dimeric kinesin is based on PDB# 4HNA (rear head in closed conformation; Gigant et al., 2013) and 4LNU (front head in open conformation; Cao et al., 2014). (**A**) The neck linker and α6 helix, which connects the neck linker, are shown in red and cyan, respectively. The I325 residue is depicted in blue spheres. In the open state, I325 residue is prohibited from binding to the complementary hydrophobic pocket due to the backward strain applied to the neck linker. In contrast, in the closed rear head, the hydrophobic interaction is stabilized by the forward strain. (**B**) ATP, P-loop, switch I and II loops in the nucleotide pocket region are shown in yellow, green, cyan and red, respectively. R203 residue in switch I and E236 residue in switch II, both essential for the hydrolysis reaction (Parke et al., 2010), and γPi of ATP are shown in space-filling. In the open head, switch I and II loops are distanced from the P-loop, whereas in the closed state, these loops approach the γPi of ATP and become capable of catalyzing the ATP hydrolysis reaction.

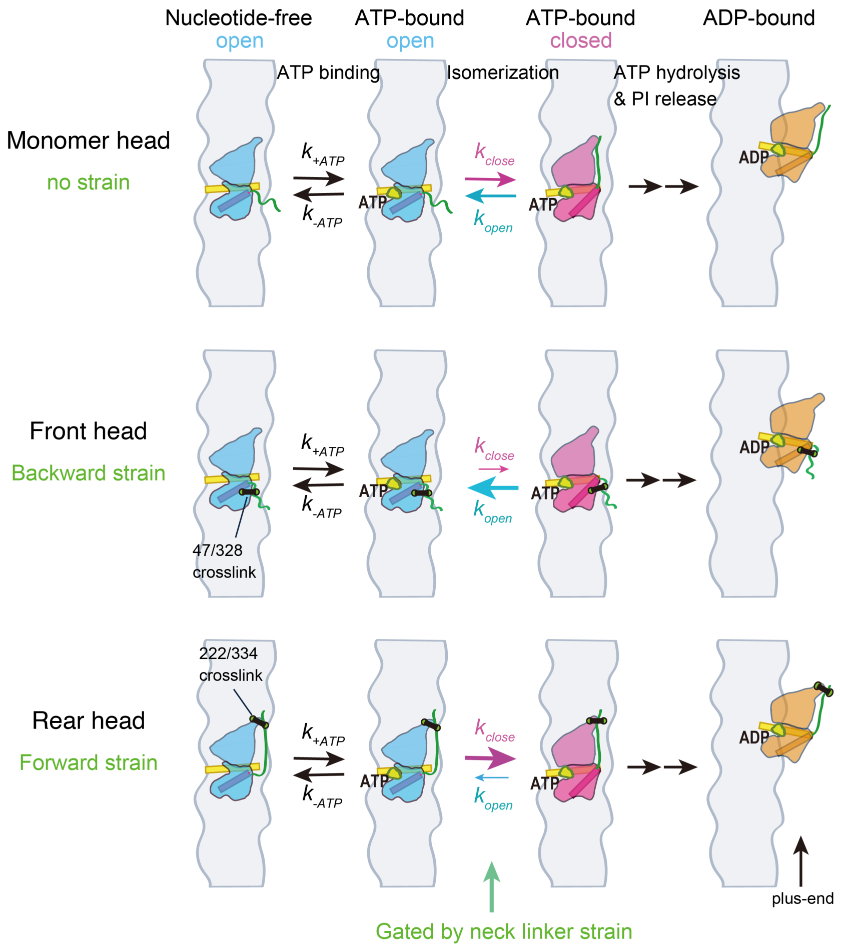


**Figure 7–figure supplement 2.** Schematic model showing that the open-closed conformational transition (isomerization) of the head is gated by the neck linker strain. The color coding is the same as in **Figure 7**. Monomer head without the neck linker strain can transit between two distinct ATP-bound configurations (**Figure 2C**; $k_{close} \sim k_{open}$): the neck-linker undocked, nucleotide-pocket opened (open; bule) state, and the neck-linker docked, nucleotide-pocket closed (closed; red) state. Backward strain applied to the front head (or C47/C328 crosslink) shifts the equilibrium towards the open state ($k_{close} < k_{open}$), suppressing ATP hydrolysis and promoting reversible ATP dissociation. On the other hand, forward strain applied to the rear head (or C222/C334 crosslink) shifts the equilibrium towards the closed state ($k_{close} > k_{open}$), increasing ATP affinity and facilitating ATP hydrolysis. We speculate that the rear head initially adopts an open conformation after ATP binding and the ATP binding kinetics to the open head ($k_{+ATP}$ and $k_{-ATP}$) is the same as the monomer without neck linker constraint. Assuming the transition rate to the closed state significantly exceeds the ATP dissociation rate in the rear head ($k_{-ATP} < k_{close}$), the apparent ATP release rates ($k_{-1}$) observed for the E236A mutants (**Figures 2D** and **5B**) are dependent on the rate constants for the transition from closed to open state ($k_{open}$). As a result, $k_{-1}$ of E236A mutants become lower as the forward strain increases (which decreases the $k_{open}$) (**Figures 5D**).
